# Tn7-CRISPR-Cas12K elements manage pathway choice using truncated repeat-spacer units to target tRNA attachment sites

**DOI:** 10.1101/2021.02.06.429022

**Authors:** Shan-Chi Hsieh, Joseph E. Peters

**Affiliations:** Department of Microbiology, Cornell University, Ithaca, NY 14853, USA

**Keywords:** CRISPR regulation, Guide RNA categorization, RNA-directed transposition

## Abstract

CRISPR-Cas systems provide a defense against mobile elements. These defense systems have been naturally coopted multiple times for guide RNA-directed transposition by Tn7-like transposons. Elements associated with a type I-F CRISPR-Cas system categorize guide RNAs, maintaining a standard CRISPR array capable of acquiring new spacers targeting other mobile elements while maintaining a special guide RNA allowing integration into a conserved site in the chromosome called an attachment site. We show here that Tn7-like elements associated with a type V-K (Cas12K-based) system use a similar strategy to target diverse tRNA genes as attachment sites. These guides are encoded as truncated minimal repeat-spacer units and are found in distinct locations. Multiple pieces of information support that V-K guide RNAs are acquired using a type I-D adaptation system, but remain private to the V-K transposition process. This catalog of Cas12K elements and naturally occurring insertions will help future work engineering precision integration systems.

## Introduction

CRISPR-Cas systems serve as adaptive immune systems that help bacteria and archaea exclude mobile elements like bacteriophages, transposons, and plasmids^1^. Intriguingly, multiple examples exist where CRISPR-Cas defense systems have been coopted by mobile elements to support their own maintenance and dispersal^2^. Of interest are multiple examples where CRISPR-Cas systems associate with a specialized type of transposons called Tn7-like elements^12, 13^. These natural systems of guide RNA-directed transposition show promise as tools for precision integration of genetic payloads, but fundamental questions remain with how they normally function and how they interact with canonical active CRISPR-Cas systems^4-7^.

All of the known associations between CRISPR-Cas systems and transposons occurred with Tn7-like elements, which have a dedicated system of target site selection^12, 13^. In addition to the transposase that is responsible for recognizing the *cis*-acting ends of the element and joining them to a new target DNA, Tn7-like elements have a conserved transposition regulator protein, TnsC, and a target site identification protein, TniQ/TnsD. Prototypic Tn7 provides the best-understood example of this type of element, encoding five Tns proteins (TnsA, TnsB, TnsC, TnsD/TniQ, and TnsE) that allow for two targeting pathways^8^ (Figure 1). One targeting pathway recognizes a conserved attachment site in the chromosome associated with the essential *glmS* gene (called *attTn7*) in bacteria. The *glmS* coding region is directly recognized by the TnsD/TniQ protein, which recruits the TnsC regulator protein, and then in turn the TnsA+TnsB transposition machinery complexed with the Tn7 end sequences. Tn7 insertions are directed into the transcriptional terminator of the *glmS* gene at a fixed distance from the sequence recognized by TnsD/TniQ. In Tn7, a second target site selection protein, TnsE, that is unrelated to TnsD/TniQ, recognizes features of DNA replication in a process that allows preferential insertion into conjugal plasmids to maximize dispersal of the element to new bacterial hosts^9-12^.

**Figure 1.**
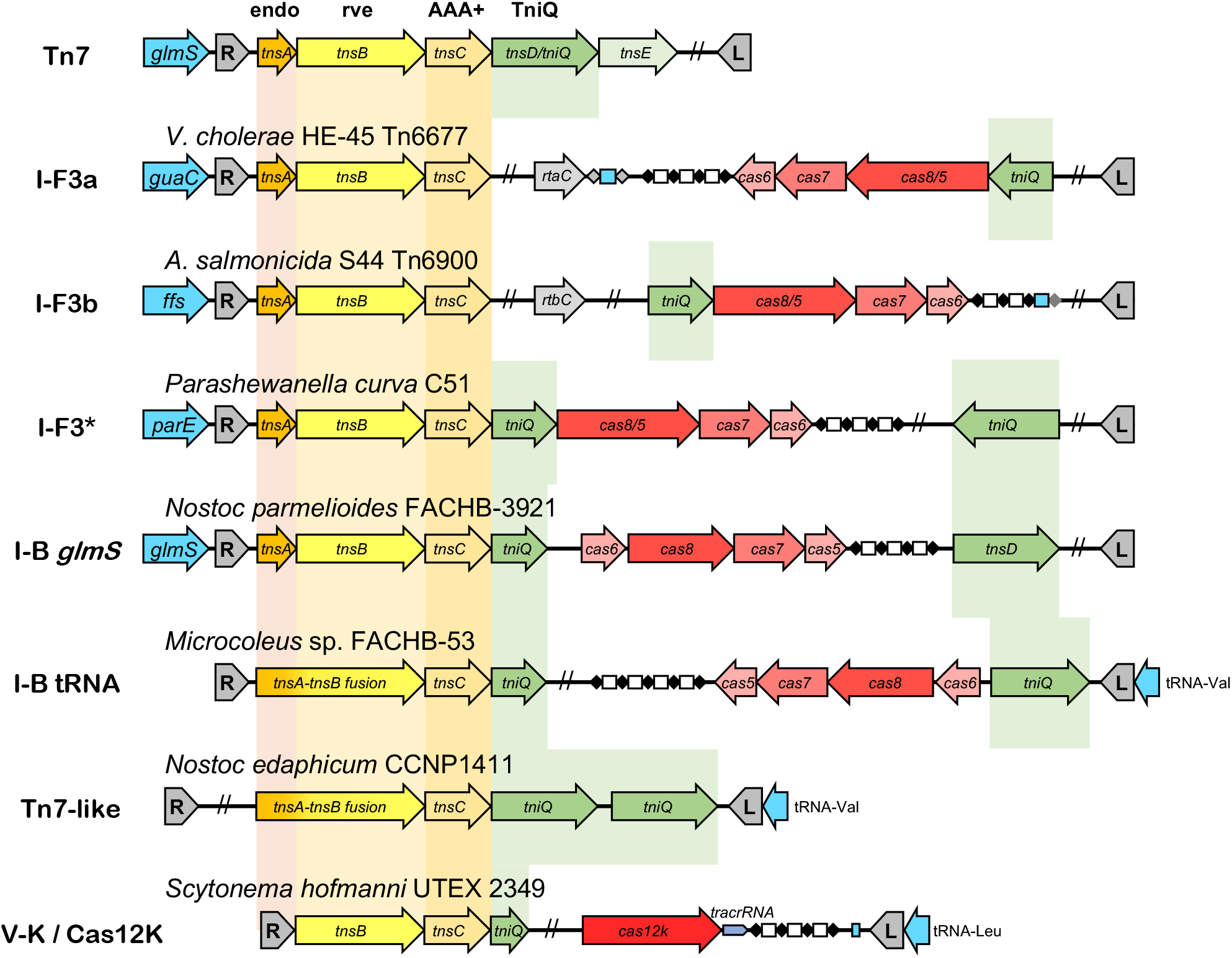
Schematic representation of selected Tn7-like elements. Tn7-like elements are distinguished by the ability to control target site selection using multiple targeting pathways. Tn7-like elements have a preferred attachment site (*att* site), typically associated with a highly conserved and essential gene (blue box arrows). In many systems a second pathway is known to allow a mechanism to target other mobile elements that can facilitate transfer of the transposon to new bacterial hosts. Prototypic Tn7 has a transposase with two proteins TnsA and TnsB, a AAA+ regulator protein called TnsC, and a target site selection protein TnsD from the TniQ-family of proteins (Box arrows). A broad diversity of Tn7-like elements shares some or all of the Tn7 core components. Derivatives have coopted CRISPR-Cas systems for targeting mobile elements as transposition targets (shades of red box arrows). These include versions that evolved from type I-F1 CRISPR-Cas systems, the I-F3a (Tn6677) and I-F3b (Tn6900) elements and a sister group indicated in the figure as I-F3^*^. A single effector Cas12K system was also coopted for guide RNA directed transposition. The CRIPSR arrays are indicated with diamonds for the repeats and rectangles for the spacers. Guide RNAs encoded in the spacers that recognize the *att*-site are in blue. The host strain for the example shown, the left (L) and right (R) ends of the transposons, known transcriptional regulators of the I-F3 systems (*rtaC* and *rtbC*), and the tracrRNA are indicated in the figure. Multiple mechanisms are used to target genes encoding tRNAs as target sites.

In one well-represented group of Tn7-CRISPR-Cas elements functionally distinct guide RNAs allow two transposition pathways analogous to Tn7^7^. These elements evolved from the canonical I-F1 CRISPR-Cas systems, as a variant called I-F3 with Cas6, Cas7, and a fused version of Cas5 and Cas8 as an effector complex^3^ (Figure 1). The type I-F3 systems lack an interference component, the Cas3 nuclease, normally found in type I systems. Like all of the Tn7-CRISPR-Cas systems, the I-F3 systems lack a Cas1Cas2 adaptation system for acquiring spacers^2, 13^. Pathway choice is regulated using either a specialized guide RNA or a guide RNA that is differentially regulated. In the I-F3 elements, a dedicated guide RNA allows long-term memory to one of a small number of conserved attachment sites recognized at the essential *ffs, guaC*, or *rsmJ* genes or a gene of unknown function, *yciA*. I-F3 systems also possess a canonical CRISPR array capable of acquiring new guide RNAs to allow access to mobile elements for dispersal of the transposons between bacteria.

A second well-represented group of Tn7-CRISPR-Cas elements, Tn7-CRISPR-Cas12K elements, use a type V, single-effector, Cas12 system^1, 2^ (Figure 1). Members of the Tn7-CRISPR-Cas12K family have been used in systems for practical applications called CAST^4^ (CRISPR-associated transposase) and INTEGRATE^13^ (Insertion of transposable elements by guide RNA-assisted targeting). Each of the subgroups of Cas12 proteins found in the type V systems (V-A through V-K) are believed to each have evolved independently from TnpB proteins from nonautonomous transposons^1, 14^. Presumably, the adaptation systems associated with each type V subgroup were independently acquired from ancestral adaptation systems. Like the I-F3 systems, the transposon-associated type V-K systems lack adaptation and interference^2, 3^. Tn7-CRISPR-Cas12K elements have TnsB, TnsC and TniQ, but lack a homolog of TnsA^2^(Figure 1). Tn7-CRISPR-Cas12K elements were previously shown to reside in the chromosome at tRNA attachment sites, but how these sites are recognized was not known^2^.

Here we show that Tn7-CRISPR-Cas12K elements use distinct dedicated guide RNAs to recognize chromosomal attachment sites. Diverse tRNA genes are primarily used as attachment sites and in a few instances other functional RNA genes or the *mutS* and *mutL* mismatch repair genes. The repeat-spacer units used for guide RNAs that recognize the chromosomal attachment sites are naturally truncated at the coding level to the previously determined minimal processed functional unit. It remains unproven how spacers are acquired with the Cas12K systems, but we propose that type I-D adaptation systems are used by the Cas12K systems based on the protospacer adjacent motif (PAM), the sequence and size of the repeat-spacer unit, and the overrepresentation of I-D systems in cyanobacteria where essentially all Tn7-CRISPR-Cas12K elements are found. Privatization of the *att*-targeting Tn7-CRISPR-Cas guide RNAs from the CRISPR-Cas system used for adaptation may occur based on a diverged repeat sequence and/or because of the minimal nature of the repeat-spacers, or an unknown mechanism. Our data provide a catalog of naturally occurring guide RNA insertions and an updated inventory of ∼150 complete Cas12K systems that should benefit future work for engineering guide RNA-based transposition systems.

## Methods

### Identifying Cas12K CRISPR-guided Tn7-like elements

Annotated protein fasta files, genomic sequences, and feature tables of 1,469 *Cyanobacteria* genomes were downloaded from National Center for Biotechnology Information (NCBI) FTP site on November 18, 2020. The Cas12K proteins were detected by sequence homology to ShCAST Cas12K with BLAST and hits 500 amino acids in length were used for further analysis^15^. Profile HMMs associated with TnsB (PF00665), TnsC (PF11426,PF05621), TniQ(PF06527) and MerR family proteins (PF13411), were downloaded from The European Bioinformatics Institute (EMBL-EBI) Pfam database and used for detecting homologs with hmmsearch (HMMER3)^16^. The TnsBCQ operons were assembled based on orientation and distance of the detected homologs and grouped with the proximal downstream *cas12k* gene. The MerR family proteins upstream of *cas12k* gene with opposite orientation were collected.

### CRISPR array detection

The CRISPR arrays were first detected with MincED (https://github.com/ctSkennerton/minced). CRISPR arrays within 3 kbp downstream of *cas12k* genes were grouped with the *cas12k* genes. These *cas12k* associated CRISPR repeats were used to construct a sequence profile, which was then used for second round CRISPR detection with nhmmscan to better define the arrays^16^.

### Attachment site targeting spacer detection

To detect attachment site targeting spacers and protospacers, the 17 bp sequences following GAAA(G/C) were collected from downstream of *cas12k* gene as candidate *att*-targeting spacers. These candidate *att*-targeting spacers were used to detect near-identical sequences (<4bp mismatches) further downstream and have orientation opposite to *cas12k*. The sequence pairs with least mismatches are collected as putative *att*-targeting spacer and protospacer for further analysis.

## Results and Discussion

### Tn7-CRISPR-Cas12K elements use guide RNAs derived from truncated spacers to target chromosome attachment sites

All of the previously identified Tn7-CRISPR-Cas12K elements were in cyanobacterial genomes^2^. For our updated analysis we downloaded 1,469 annotated cyanobacterial genomes and identified 282 relatively intact *cas12k* genes (encoding a protein 500 a.a.) based on homology to the previously identified proteins^2^ (Supplemental Table S1). In this pool we identified 152 *cas12k* genes which had a proximal *tnsBCQ* operon (Supplemental Table S1). We suggest loss of the *tnsBCQ* operon results from inactivation and subsequent degradation of elements (and not use of the Cas12K protein for other functions) because in almost all cases we could identify a candidate transposon end-sequence, a CRISPR array, and a candidate tRNA attachment site associated with the *cas12K* genes (see below). Genes for *cas12K* without an identifiable proximal *tnsBCQ* genes also encoded Cas12K proteins that did not branch separately from the other element-associated proteins, consistent with a function in transposition (Figure 2).

**Figure 2.**
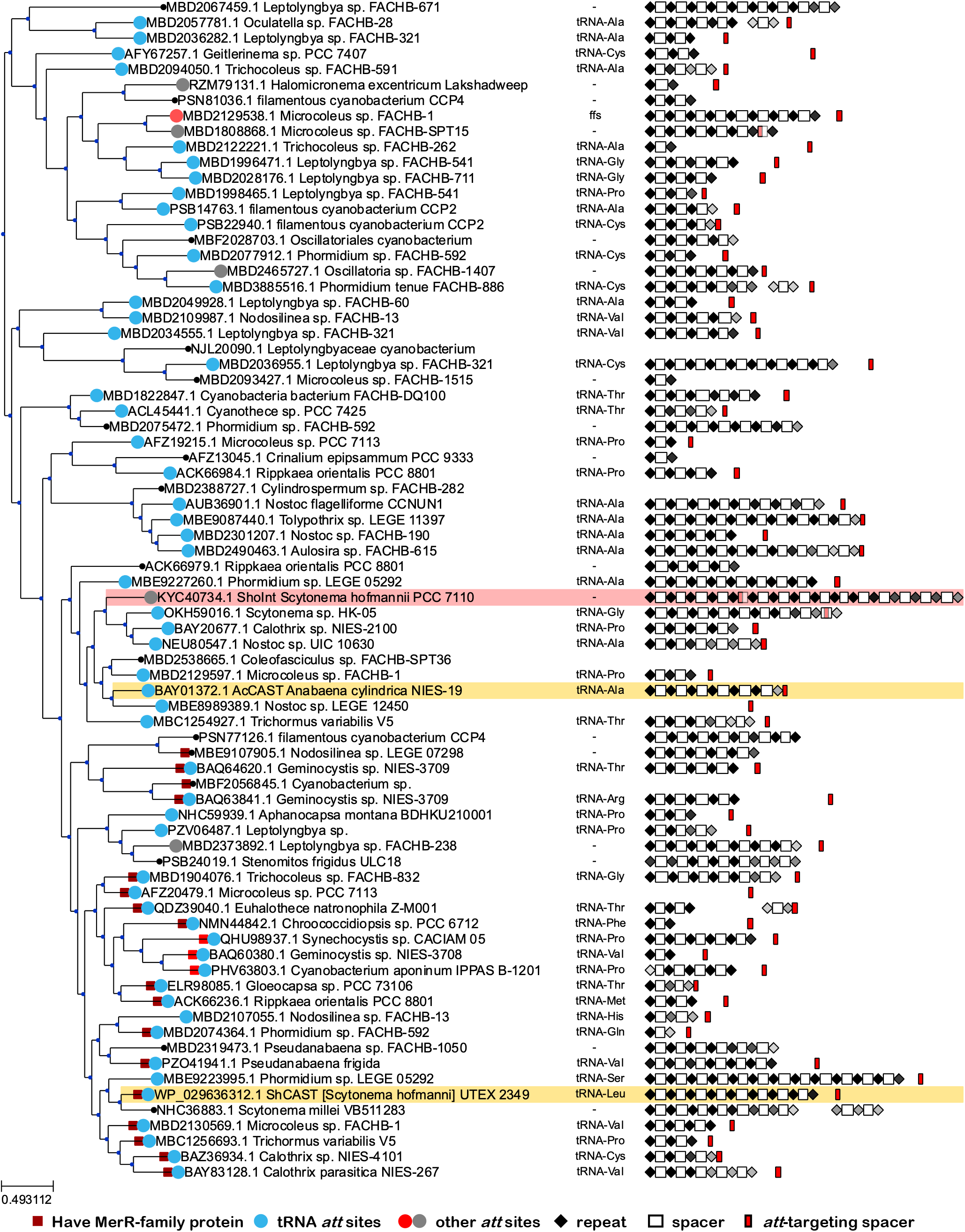
Cas12K similarity tree with *att* sites and the CRISPR array configuration with the relative position of the truncated repeat-spacer unit. A Cas12K similarity tree representing 252 elements indicated by host strain. Elements that were more than 70% identical are represented in the figure with a single representative. The attachment site was a tRNA gene in almost all cases as indicated. CRISPR arrays are indicated with diamonds for the repeats and rectangles for the spacers. More diverged repeats are indicated in grey. Guide RNAs encoded as truncated repeat-spacer units that recognize the chromosomal attachment site are indicated in red. Systems previously established in a heterologous *E. coli* host are indicated with yellow^4^ and pink^13^ highlight. Other features are indicated in the figure. See text for details.

As part of our bioinformatics analysis, we searched for candidate attachment-site-targeting (*att*-targeting) guide RNAs encoded within the transposon. In our pool of 282 elements, CRISPR arrays had an average repeat length of 37 bp and a spacer length of 36 bp and varied from one to fifteen guides. Notably, in essentially all cases we could also identify a candidate 17 bp spacer as part of an *att*-targeting guide RNA in the non-coding region downstream of *cas12k* gene (Figures 1-2 and Supplemental Figure S1). These were missed in the original description of these elements because of differences from the standard repeat-spacer configuration found in the array. The candidate *att*-targeting guide RNAs invariably possess a ∼13bp sequence resembling the 3’ end of associated CRISPR repeats (Figure 3). Interestingly, the 17 bp spacer sequences and their upstream ∼13bp repeat sequences correspond to the size of the processed crRNA unit revealed by previous work with RNA sequencing^4^. We could not detect any conserved sequence motifs outside the putative crRNA forming unit, implying the sequence alone may be sufficient for crRNA processing and RNA-guided transposition into chromosome attachment sites (Figure 3). Because the sequences usually only contain partial CRISPR repeat and 17 bp spacer, we refer to them as truncated repeat-spacer units hereafter.

**Figure 3.**
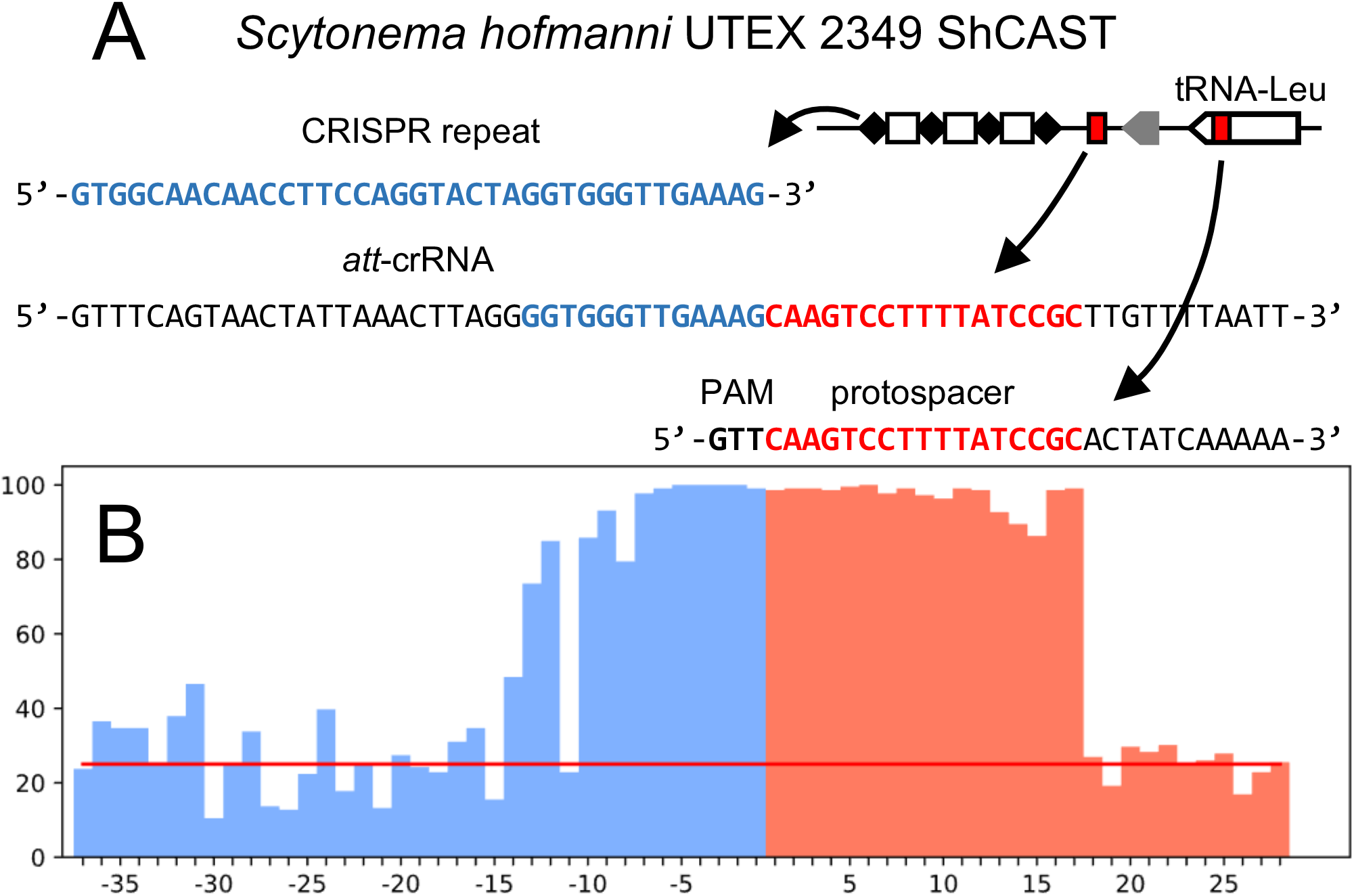
Representative truncated repeat-spacer units. The Cas12K element guide RNAs are encoded in CRISPR arrays with 37 bp repeats and ∼35 bp spacers. Guide RNAs recognizing chromosomal attachment sites are truncated relative to the CRISPR array configuration. (A) A representative CRISPR repeat from *Scytonema hofmanni* UTEX 2349 ShCAST and the attachment site targeting truncated repeat-spacer unit (*att*-crRNA) are indicated relative to the position in the element. The *att*-crRNA encoding sequence only contains the tracrRNA base-pairing region of the CRISPR repeats, and only matches 17bp of the protospacer. (B) The match between each position of *att*-targeting crRNA encoding region and CRISPR repeats (blue) or protospacers (red) from 219 Cas12K elements with *att*-targeting crRNA is indicated as a percentage. Red line marks the 25% line. See text for details.

The *att*-targeting guide RNAs were typically encoded at the end of the CRISPR array or further 3’ to the array (Figure 2 and Supplemental Table S1). Additional features also support that a guide RNA derived from the truncated repeat-spacer unit was responsible for targeting the chromosome attachment sites used by the elements. The corresponding protospacers are an exact match to the size of the spacer (17 bp) and in most cases are a perfect sequence match (Supplemental Table S1). Additionally, nearly all of the protospacers have a GTN protospacer adjacent motif (PAM) (Supplemental Figure S1), which confirms the PAM screening results of previous study^4^ but in the natural setting. The orientation of the elements and the spacing from the protospacer was also consistent with previous results found with the system in *E. coli*^4, 13^ (Supplemental Figure S1).

### Dual pathway functionality in Tn7-CRISPR-Cas elements occurs either by guide RNA categorization or the use of multiple TniQ proteins

Our identification of distinct, *att*-targeting guide RNAs in Cas12K elements is similar to our previous finding with I-F3 elements^7^. Similar to the I-F3 elements, Cas12K elements possess *att*-targeting spacers, and the locations of *att*-targeting spacers are either at the end of CRISPR array or further downstream (Figure 2). Such an arrangement allows the transposons to have two transposition pathways with the same protein components. Our finding that a system of categorizing RNAs evolved independently twice suggests there are important benefits to this form of target site selection.

The I-F3 elements that show guide RNA categorization, also have a sister group that uses two TniQ proteins to allow the use of dual pathways; one that is dedicated to a *parE* chromosomal attachment site recognized by TniQ and the second functioning as a dedicated part of the I-F3 CRISPR-Cas system^7^(Figure 1). Two independently evolved families of type I-B Tn7-CRISPR-Cas elements also have two TniQ proteins in each element^3, 6^ and seem likely to share a similar division of functions (Figure 1).

### Diverse tRNAs are used as *att* sites suggesting a special advantage as a preferred integration site

Compared with I-F3 elements, the chromosome attachment sites of Cas12K Tn7-CRISPR-Cas elements are much more diverse. Our previous work found 94% of the genetically diverse I-F3 elements reside in one of only four different chromosomal *att* sites; Cas12K elements on the other hand are associated with at least 15 different tRNA genes, *mutS, mutL, ffs* (Signal recognition particle RNA gene), a nuclease, and a putative sRNA (Supplemental Table S1). Unlike I-F3 elements, the attachment sites of Cas12K elements don’t follow the phylogeny of the *cas12K* genes encoded by the elements. Even when the same type of tRNA is recognized in many cases it is via a protospacer at a different position of the gene (Supplemental Figure S1). We suspect that the bias of these elements to use diverse tRNA genes stems from a strong selective advantage in using tRNAs as attachments sites and not necessarily a bias in spacer acquisition from tRNA encoding genes. Genes for tRNAs are known to be hotspots for attachment sites for diverse genetic elements including bacteriophages, self-splicing introns, and integrating conjugating elements^17-19^. Even within cyanobacteria, another unrelated Tn7-like element is known to target tRNA genes likely by direct recognition of the coding region of the gene (Figures 1 and 4). These elements were recognized previously and have a fused version of the TnsA and TnsB proteins^3, 6^. The TniQ protein associated with tRNA targeting at the DNA level have Xre and AcrR-type DNA binding domains^20^. The tRNA-genes used as *att* sites appear to be multiply represented in cyanobacteria^21^, which might also provide an advantage to the transposon. It was previously suggested that integration downstream from a tRNA gene may allow the element to benefit from the expression profile of the tRNA genes^18^, a similar benefit was argued for Tn7 elements that integrate downstream of the *glmS* gene^22^.

**Figure 4.**
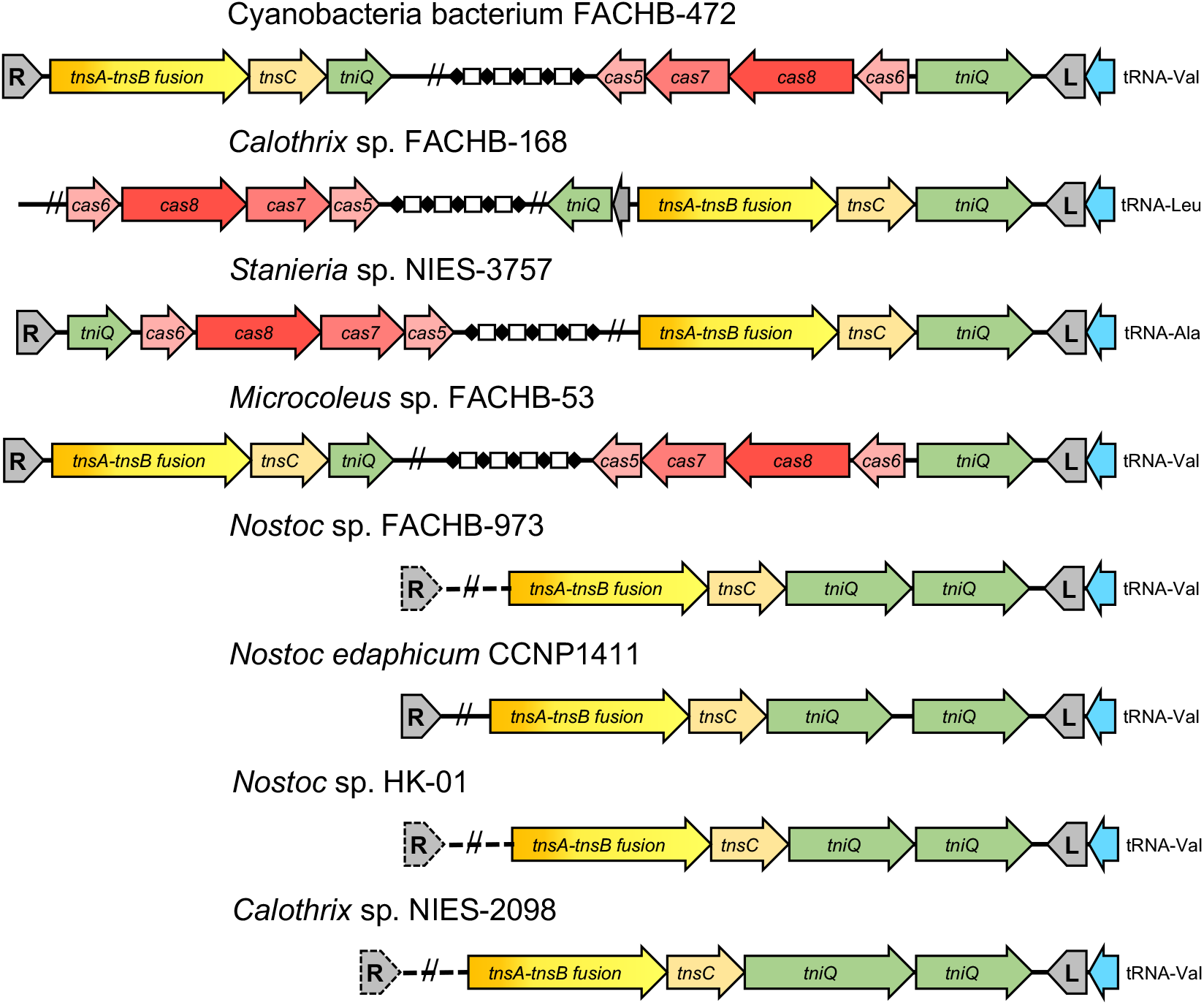
Tn7-like elements targeting tRNA genes in cyanobacteria. Representative Tn7-like elements from cyanobacteria targeting tRNA genes. Unlike Tn7 and other Tn7-like elements the attachment site was adjacent to the end of the element that was not proximal to the transposase genes (The right end is operationally defined as the end next to the transposase gene(s)). TnsA and TnsB fused in all of the systems shown. Features of the figure are as described in Figure 1. In one case the candidate right end (R) could not be identified, in some cases the candidate right end of the element was situation elsewhere in the chromosome (indicated with a dashed line).

Beyond tRNA genes, other *att*-sites found with the Cas12K elements include functional RNAs, *ffs* and another putative sRNA. Both tRNA and *ffs* genes are expected to be highly conserved and functional RNA genes may be more conserved than protein coding genes that show more variability from the third position in codons. Having a perfect or near perfect match may be more important with the 17 bp protospacer found with Tn7-CRISPR-Cas12K elements driving a bias to functional RNA genes. It is unclear if there is any significance to the finding that both components of the mismatch repair system, *mutS* and *mutL*, serve as *att* sites recognized by Tn7-CRISPR-Cas12K elements.

### Cas12K elements may acquire spacers using type I-D CRISPR-Cas adaptation systems

Like other guide RNA-directed transposons, the Cas12K elements lack adaptation genes and therefore must rely on other CRISPR systems for spacer acquisition. However, unlike I-F3 elements and I-B elements, which have their canonical versions of CRISPR systems as potential helpers, CRISPR-Cas12K and other type V systems are believed to have evolved independently^1,14^. Consequently, Cas12K elements must rely on another CRISPR-Cas system of a different type for adaptation. A similar scenario is found with type III and type IV CRISPR-Cas systems which are rarely associated with adaptation systems^1^.

We suggest that Cas12K systems use the Cas1Cas2Cas4 adaptation system from type I-D systems for acquiring spacers. Type I-D CRISPR-Cas systems provide a good initial candidate as they are one of the most abundant systems in cyanobacteria^23^, where nearly all of the Tn7-CRISPR-Cas12K elements are found. The currently characterized type I-D CRISPR systems are also known to use the same GTN PAM found in Cas12K systems^24, 25^ (Supplemental Figure S1 and Supplemental Table S1). Furthermore, the length of type I-D CRISPR repeats is 37 bp, identical to the repeat length we identify in most of the Tn7-CRISPR-Cas12K element arrays (Supplemental Table S1). This is in spite of the finding that the sequence transcribed from the spacers is processed down to 17 bp when matured into a Cas12K effector complex^4^. The actual repeat sequence found in I-D and V-K systems also show intriguing similarities. While the sequence of the repeats found in Cas12K CRISPR arrays is highly variable, they do show conservation at the 5’ and 3’ ends of the repeats (Supplemental Figure S2). Interestingly, this same sequence is found conserved with the I-D CRISPR array repeats (Figure 5). The spacer length distribution in type I-D CRISPR is also similar to Cas12K, averaging around 35 bp and varying by 5 bp. Finally, a sequence found conserved across the leader region in the arrays associated with Cas12K elements matches the sequence and position (∼-44 bp) of a motif important for spacer acquisition in I-D systems (Figure 5). Although direct experimental validation would be needed to confirm what seems apparent from our bioinformatic analyses, it appears likely that Tn7-CRISPR-Cas12K arrays rely on cross-class form of adaptation for acquiring spacers. If the above model is correct, it would also suggest a mechanism that could allow guide RNAs encoded in the Cas12K array to remain private to the transposon system. The self-targeting guide RNAs used to recognize attachment sites are in truncated repeat-spacer units which lack the sequences needed for use in a type I-D effector complex^25^. Even the guide RNAs encoded in the standard Cas12K CRISPR array are likely private to the transposon, given they lack the stem-loop used for Cas6 processing by the type I-D systems^26^.

**Figure 5.**
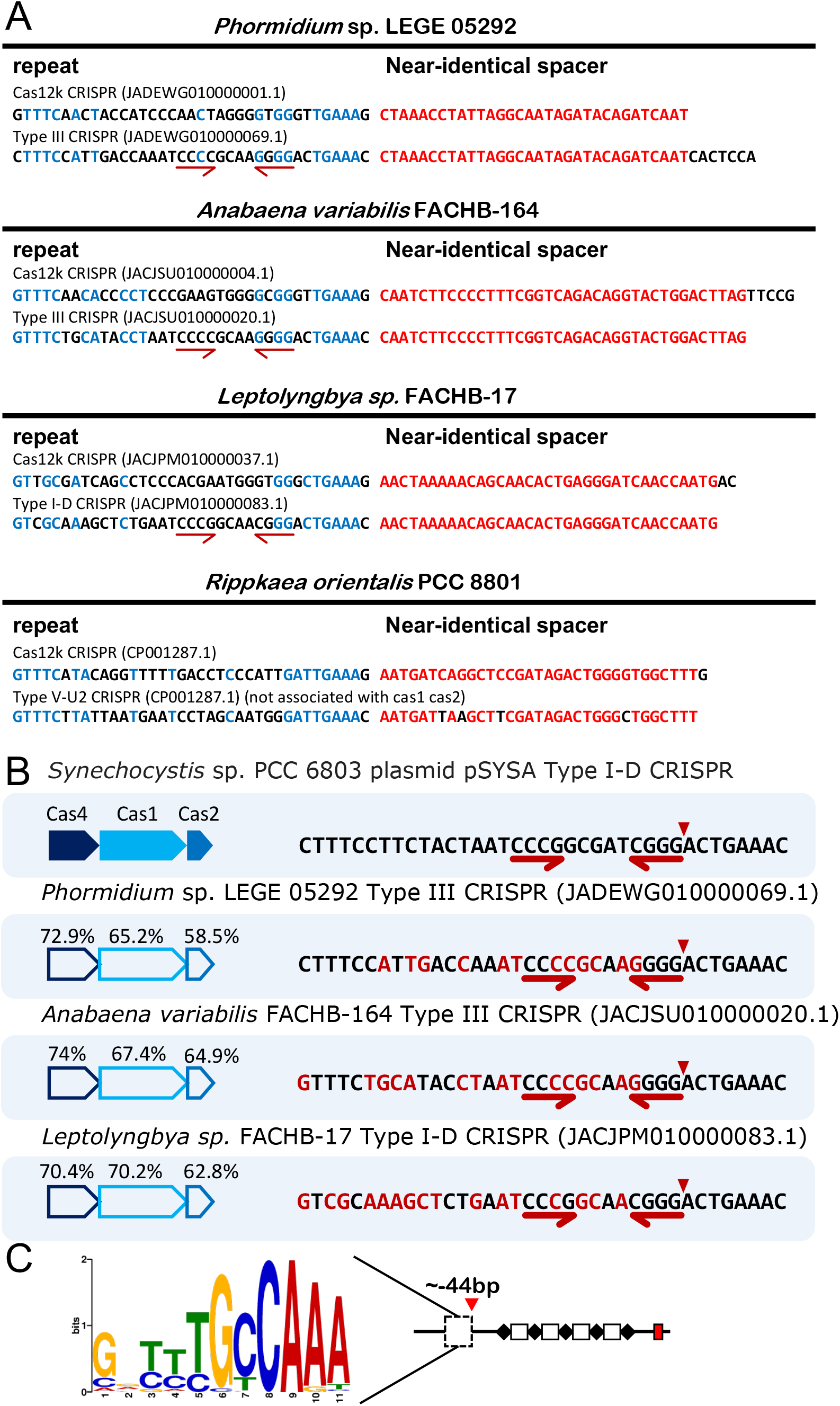
Near-identical spacers across CRISPR arrays associated with diverse CRISPR-Cas systems related to features of type I-D adaptation systems. Four cases where there were near-identical spacers both in a Cas12K associated array and an array associated with another type of system within the same genome. (A) Near-identical spacers identified in Cas12K associated arrays were identified in type III, type I-D, and type V-U2 systems in cyanobacteria. Blue typeface indicates conserved positions in the repeats and red typeface conserved position in the spacers. The GenBank accession number is indicated in parenthesis. (B) Homology between a representative Cas1Cas2Cas4 I-D adaptation system from cyanobacteria and system shown in panel A. Conserved features of the of the leader sequence are indicated with half arrows and the relative point of Cas4 processing is indicated with a triangle. (C) WebLogo of a conserved feature found in the leader region of Cas12K CRISPR arrays and in type I-D CRISPR-Cas systems^34^.

Interestingly, in the course of our work we found four cases where there were near identical spacers both in a Cas12K associated array and an array associated with another type of system within the same genome (Figure 5). These include arrays associated with two type III systems, a type I-D system, and a type V-U2 system. This could be consistent with the same adaptation functioning across all of these CRISPR-Cas systems. The CRISPR array repeats were all 37 bp and showed the same conservation with Cas12K repeats at the 5’ and 3’ ends (Supplemental Figure S2). While it is an enticing idea, additional work will be needed to determine if type I-D adaptation systems are special in their ability to function across multiple classes and types of CRISPR-Cas systems.

### Regulation of pathway choice with Tn7-CRISPR-Cas12K elements

It is unclear if Tn7-CRISPR-Cas12K elements have a mechanism to actively regulate pathway choice. One branch of Tn7-like elements with the type I-F3 CRISPR-Cas system use an Xre-family protein for a form of zygotic induction to strongly induce the expression of a guide RNA specific to the chromosomal attachment site when they enter a new host (*rtaC* in Figure 1). The *att*-targeting guide RNA is subsequently strongly repressed once the element is established in the host^7^. No known transcriptional regulators were found broadly conserved across Tn7-CRISPR-Cas12K elements in our analysis. While we did identify a MerR-family DNA binding protein conserved in one branch of the Cas12K elements, there were no obvious DNA binding motifs that would suggest they are responsible for regulating access to chromosomal attachment sites (Figure 2 and Supplemental Table S1).

### DNA processing events with Tn7-CRISPR-Cas12K elements

A mystery that remains with Tn7-CRISPR-Cas12K elements involves details of the post transposition processing events. Tn7-CRISPR-Cas12K elements lack the TnsA protein normally associated with Tn7-like elements^2,6,27^(Figure 1). TnsA is an endonuclease that is responsible for allowing Tn7-like elements to move using a cut-and-paste mechanism, where the element if fully removed from the donor position during transposition^28-30^. Transposons, like Cas12K elements, with homologs of the TnsB, TnsC, and TnsD/TniQ protein move using a replicative transposition mechanism, which forms a co-integrate between the donor and target DNAs ^31^. These families of transposons encode a second recombination system, a site-specific recombination system that allows for resolution of cointegrates. We failed to observe conserved site-specific recombination systems in our pool of Cas12K elements that could resolve cointegrates in these systems. It therefore remains unclear how processing proceeds in the natural setting. While cointegrate events are found with the Cas12K systems when transposition was established in *E. coli*^32^ it is unclear what occurs in cyanobacteria where the element is normally found. Cointegrates between plasmids and bacteriophage that served as the donor for the Cas12K element are not expected to be tolerated if integrated into the chromosome. Among the possibilities are that a specialized DNA processing activity is found natively in cyanobacteria that serves the same biochemical function of TnsA or that a recombination system found in cyanobacteria is particularly efficient at resolving cointegrates. Cyanobacteria are often polyploid and have previously been reported to be genetically unstable, which could be consistent with homologous recombination systems with unique properties^33^.

## Conclusion

Tn7-CRISPR-Cas12K elements use a form of guide RNA categorization possessing a standard CRISPR array in addition to encoding a truncated repeat-spacer unit for an *att*-targeting guide RNA. The *att*-sites recognized are typically diverse tRNA genes. Based on similarities between the CRISPR arrays, Type I-D CRISPR-Cas systems may be responsible for capturing spacers in the Tn7-CRISPR-Cas12K systems. Our data provides a series of candidates to help develop precision integration of genetic payloads using guide RNA-directed transposition.

## Acknowledgements

We thank Zachary Barth and Michael Petassi for comments on the manuscript.

## Author Confirmation Statement

SCH carried out the bioinformatics analyses with assistance from JEP. SCH and JEP wrote the paper. Both authors have reviewed and approve of the manuscript. This manuscript was submitted solely to this journal and is not published, in press, or submitted elsewhere.

## Author Disclosure Statement

The lab has corporate funding for research that is not directly related to work in this publication.

## Funding Statement

This work was supported by NIH R01 GM129118 and R21 AI148941.

## Supplemental figure titles and legends

**Supplemental Table S1 – Cas12K associated elements identified in the study**

**Supplemental Figure S1 – Cas12K similarity tree with the PAM and the relative position of the protospacer in the target gene**

A Cas12K similarity tree representing 252 elements indicated by host strain. Elements that were more than 70% identical are represented in the figure with a single representative. The attachment site was a tRNA gene in almost all cases and the relative position of the protospacer in the target gene indicated in red. The three base pair PAM and its spacing from the left end of the element is indicated. Other features are indicated or as described in the Figure 2 legend.

**Supplemental Figure S2 – Cas12K similarity tree with WebLogos of the repeats**

A Cas12K similarity tree representing 252 elements indicated by host strain. Elements that were more than 70% identical are represented in the figure with a single representative. Figure shows the WebLogo for the repeats in the element CRISPR array. Other features are indicated or as described in the Figure 2 legend. See text for details.

