## Supplementary figures and images for "Tn7-CRISPR-Cas12K elements manage pathway choice using truncated repeat-spacer units to target tRNA attachment sites"

### Supplemental Figure 1

#### Slide 1
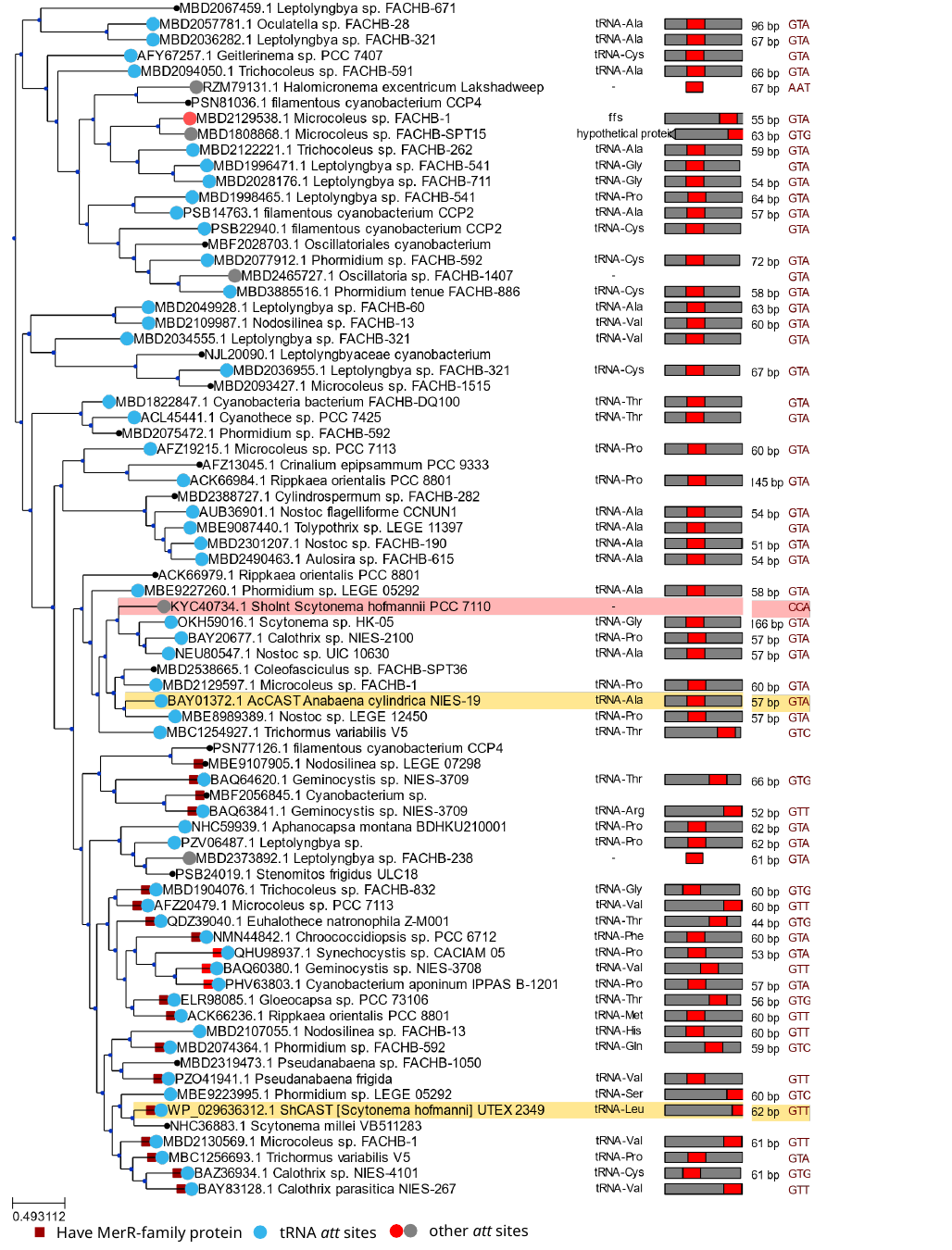

other att sites
Have MerR-family protein
tRNA att sites

### Supplemental Figure 2

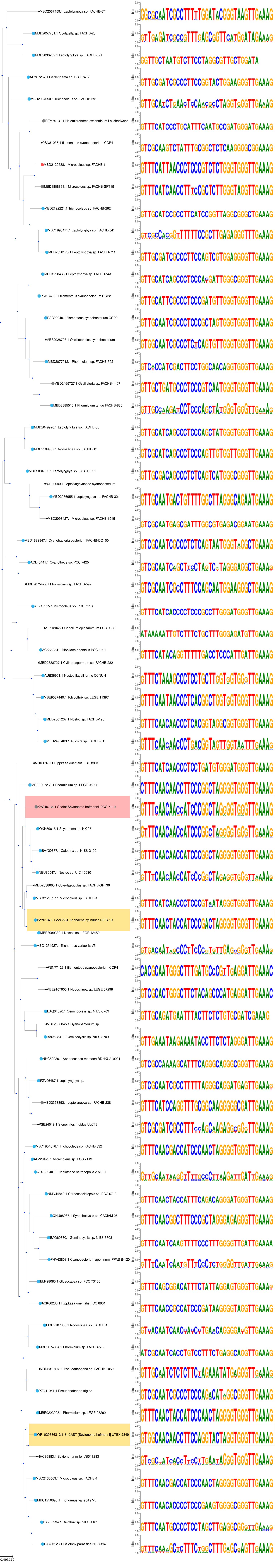
